# Viral-mediated delivery of morphogenic regulators enables leaf transformation in *Sorghum bicolor* (L.)

**DOI:** 10.1101/2025.02.15.637725

**Authors:** Nathaniel M. Butler, Aidan T. Carlson, Colby Starker, Daniel F. Voytas

## Abstract

Recent advancements in monocot transformation, using leaf tissue as explant material, have expanded the number of grass species capable of transgenesis. However, the complexity of vectors and reliance on inducible excision of essential morphogenic regulators have so far limited widespread application. Plant RNA viruses, such as Foxtail Mosaic Virus (FoMV), present a unique opportunity to express morphogenic regulator genes, such as *Babyboom* (*Bbm*), *Wuschel2* (*Wus2*), *Wuschel-like homeobox protein 2a* (*Wox2a*), and the GROWTH-REGULATING FACTOR 4 (GRF4) GRF-INTERACTING FACTOR 1 (GIF1) fusion protein transiently in leaf explant tissues. Furthermore, altruistic delivery of conventional and viral vectors could provide opportunities to simplify vectors used for leaf transformation— facilitating vector optimization and reducing reliance on morphogenic regulator gene integration. In this study, both viral and conventional T-DNA vectors were tested for their ability to promote the formation of embryonic calli, a critical step in leaf transformation protocols, using *Sorghum bicolor* leaf explants. Although conventional leaf transformation vectors yielded viable embryonic calli (43.2 ± 2.9%: GRF4-GIF1, 50.2± 3%: *Bbm*/*Wus2*), altruistic conventional vectors employing the GRF4-GIF1 morphogenic regulator resulted in improved efficiencies (61.3 ± 4.7%). Altruistic delivery was further enhanced with the use of viral vectors employing both GRF4-GIF1 and *Bbm*/*Wus2* regulators, resulting in 75.1 ± 2.3% and 79.2 ± 2.5% embryonic calli formation, respectively. Embryonic calli generated from both conventional and viral vectors produced shoots expressing fluorescent reporters, which were confirmed using molecular analysis. This work provides an important proof-of-concept for use of both altruistic vectors and viral-expressed morphogenic regulators for improving plant transformation.

## Introduction

Most crop production worldwide is confined to three monocot species: wheat, rice and maize. The growing world population, increased demand for biofuels, and changes in the climate have intensified the need for more productive varieties and their cultivation on suboptimal land (FAO, 2023). In the last half century, genetic engineering has supplemented traditional breeding efforts to keep pace with these challenges (Altpeter *et al*., 2016). This is particularly the case with the advent of genome editing technologies (Butler *et al*., 2017; Cardi *et al*., 2023). Despite these advancements, implementation of genetic engineering technologies in monocot species has been lagging, largely due to recalcitrance of monocot species to infection by *Agrobacterium tumefaciens* and reliance on specialized explant tissues and genotypes for gene delivery and regeneration (Youngstrom *et al*., 2024; Parrott *et al*., 2024).

A breakthrough in monocot transformation technology came with the use of morphogenic regulators, *Baby boom* (*Bbm*) and *Wuschel2* (*Wus2*), which greatly extended the number of genotypes capable of genetic transformation and created a new opportunity for innovation (Lowe *et al*., 2016; Yavuz *et al*., 2020; Aesaert *et al*., 2022; Zhou *et al*., 2022). Despite the improvement in transformation efficiencies, most monocot protocols utilizing growth regulators, which later included the GROWTH-REGULATING FACTOR 4 (GRF4) GRF-INTERACTING FACTOR 1 (GIF1) fusion protein, still rely on immature embryos or embryonic calli for creation of new genetically engineered events.

Recently, the authors that published initial reports using morphogenic regulators have demonstrated improvements in the use of readily available leaf tissue for genetic transformation and regeneration in a variety of monocot species using morphogenic regulators with optimized expression (Wang *et al*., 2023). The so-called leaf-base transformation protocol allows for high-throughput explant preparation using a food-processor, but in line with previous protocols using morphogenic regulators, requires genetic excision of regulator genes to recover engineered events. Therefore, transient or inducible expression of morphogenic regulators may further improve leaf-base transformation protocols.

Plant RNA viruses, such as Foxtail Mosaic Virus (FoMV), have become powerful biotechnological tools for transient gene expression (Mei *et al*., 2019), gene silencing (Liu *et al*., 2016; Mei *et al*., 2016), genome editing (Baysal *et al*., 2024a; Beernink *et al*., 2022), and they have the potential to further enhance existing genetic engineering protocols in monocot species (Scholthof and Scholthof, 2023; Abrahamian *et al*., 2020). FoMV is a positive-strand, monopartite RNA virus and member of the *Potexvirus* genus. Potexviruses, such as Potato Virus X (PVX), are highly infectious across related host species and typically cause mild symptoms such as dwarfism, mosaic patterns of necrosis, and ringspots (Whitham *et al*., 2007; Abrahamian *et al*., 2020). Biotechnological adaptations of Potexviruses have enabled transient expression of proteins and RNA without relying on introduction of DNA (Bedoya *et al*., 2012; Bedoya *et al*., 2010; Ibrahim *et al*., 2019; Khakhar and Voytas, 2021).

In this study, morphogenic regulators – maize *Bbm* (*Bbm*), *Wus2* (*Wus2*), *Wox2a* (*Wox2a*) and wheat GRF4-GIF1 (GRF4-GIF1) – were cloned into FoMV and tested as viral altruistic vectors using the Wang *et al*. 2023 leaf-base transformation protocol as a proof-of-concept in sorghum. Experiments incorporating morphogenic regulators on FoMV and conventional T-DNA vectors demonstrated they were capable of generating transgenic embryonic calli and shoots, but not in the absence of the regulators. *Bbm*/*Wus2* and GRF4-GIF1 viral vectors outperformed conventional, altruistic T-DNA vectors. Furthermore, detection of FoMV in embryonic calli was drastically reduced two-weeks post infection. This work demonstrates a new application of plant RNA viruses for enabling monocot transformation and may be applied to other plant transformation protocols.

## Results

### Traditional T-DNA vectors for sorghum leaf-base transformation

Previous reports using leaf-base explant material for monocot transformation have relied on expression of the maize *Bbm* and *Wus2* morphogenic regulators using constitutive promoters in specific vector configurations, including elements for recombinase excision (Wang *et al*., 2023). These traditional T-DNA vectors were recreated using the MoClo cloning system for both *Bbm*/*Wus2* (C-BBMWUS) and wheat GRF4-GIF1 (C-GRFGIF) regulators, with the rice *Elongation Factor 1a* (*OsEf1a*) promoter expressing the GRF4-GIF1 fusion protein (**Figure 1A, 1C**). The maize *Ubiquitin* promoter (*ZmUbi*) was used to express both *Bbm* and *Wus2* separately with *Bbm* additionally driven by a 3x enhancer. The rice *Ubiquitin* promoter (*OsUbi*) was used to drive expression of the AmCyan fluorescent marker for visual detection of transgenic calli and shoots (**Figure 1A, C, D**).

**Figure 1.**
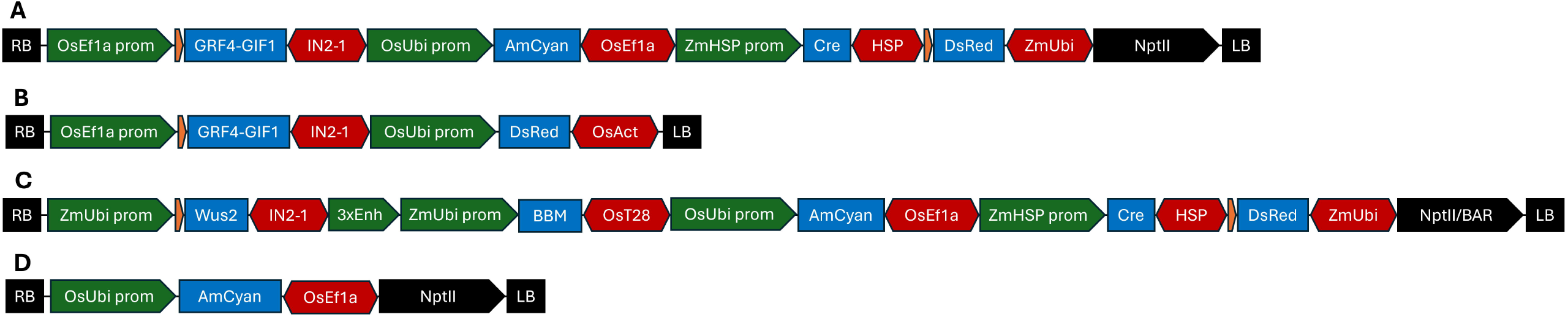
Traditional T-DNAs used for sorghum leaf-base transformation. Excisable (**A** [C-GRFGIF], **C** [C-BBMWUS]), altruistic (**B** [A-GRFGIF]), and target (**D** [T-AmCyan]) vectors for conventional and virus-mediated transformation expressing wheat GRF4-GIF1 (**A, B**), maize *Bbm*/*Wus2* (**C**) morphogenic regulators, or no morphogenic regulators (**D**). Green arrows indicate promoter or enhancer elements. Blue boxes indicate coding sequences. Red hexagons indicate terminator elements. Orange arrows indicate lox 71 (left) and lox 66 (right) sites. Black boxes indicate right and left T-DNA borders. Black arrows indicate selection marker cassettes driven by the *PvUbi2* promoter and *OsAct2* terminator. Genetic elements and assembly methods from Chamness *et al*., 2023 for incorporation into the pMIN-Ri binary vector backbone.

### Sorghum leaf-base transformation using traditional T-DNA vectors

Traditional T-DNA vectors were tested using the Wang *et al*., 2023 leaf-base transformation protocol in sorghum to determine base-line efficiencies for generating viable and transgenic embryonic calli (**Figure 2**). Control vectors containing only AmCyan (T-AmCyan) did not result in embryonic calli formation with only explant necrosis observed (**Figure S1**). However, traditional T-DNA vectors carrying morphogenic regulators resulted in embryonic calli production with calli surviving beyond four-weeks post infection and expressing AmCyan (**Figure 2**). Percentages of embryonic calli survival were calculated with slightly higher average survival rates using C-BBMWUS (50.2 ± 3.0%) compared to C-GRFGIF (43.2 ± 2.9%). Nevertheless, more drastic differences were observed among surviving calli positive for AmCyan, with C-BBMWUS outperforming C-GRFGIF by an average of 51% (**Table 1**). This data agrees with previous reports comparing the morphogenic regulators using immature embryos and provides a baseline for testing using leaf-base explants (Youngstrom *et al*., 2024).

**Table 1.** Transgenic embryonic calli efficiency using sorghum leaf transformation and conventional T-DNA vectors. Data was collected per plate with four seedlings used for each plate (“plate #”). Total embryonic calli was counted two weeks post infection (“total calli”). Calli survival was assessed by counting calli capable of growth at four weeks post infection (“total living calli”). Calli transgenesis was assessed using fluorescent AmCyan imaging (**Figure 2**), with positive (“# of calli with AmCyan”) calli counted two weeks post infection. Calli survival was calculated as a percentage (“% calli survival”) of “total living calli” divided by “total calli.” Calli transgenesis was calculated as a percentage (“% calli with AmCyan”) by dividing the number of AmCyan positive calli by “total living calli.” Average percentages of five biological replicate plates are provided with standard errors.

| <u>Vector</u> | <u>Plate #</u> | <u>Total calli</u> | Total living calli | <u>% calli survival</u> | # calli with AmCyan | <u>% calli with AmCyan</u> |
| --- | --- | --- | --- | --- | --- | --- |
| **C-GRFGIF** | **1** | 7 | 3 | 42.9% | 1 | 33.3% |
|  | **2** | 18 | 8 | 44.4% | 0 | 0.0% |
|  | **3** | 13 | 5 | 38.5% | 0 | 0.0% |
|  | **4** | 28 | 13 | 46.4% | 3 | 23.1% |
|  | **5** | 32 | 14 | 43.8% | 4 | 28.6% |
| **Average** |  |  |  | **43.2% ± 2.9** |  | **17.0% ± 15.9** |
| **C-BBMWUS** | **1** | 15 | 7 | 46.7% | 7 | 100.0% |
|  | **2** | 19 | 9 | 47.4% | 4 | 44.4% |
|  | **3** | 23 | 12 | 52.2% | 8 | 66.7% |
|  | **4** | 30 | 16 | 53.3% | 11 | 68.8% |
|  | **5** | 39 | 20 | 51.3% | 12 | 60.0% |
| **Average** |  |  |  | **50.2% ± 3.0** |  | **68.0% ± 20.3** |

**Figure 2.**
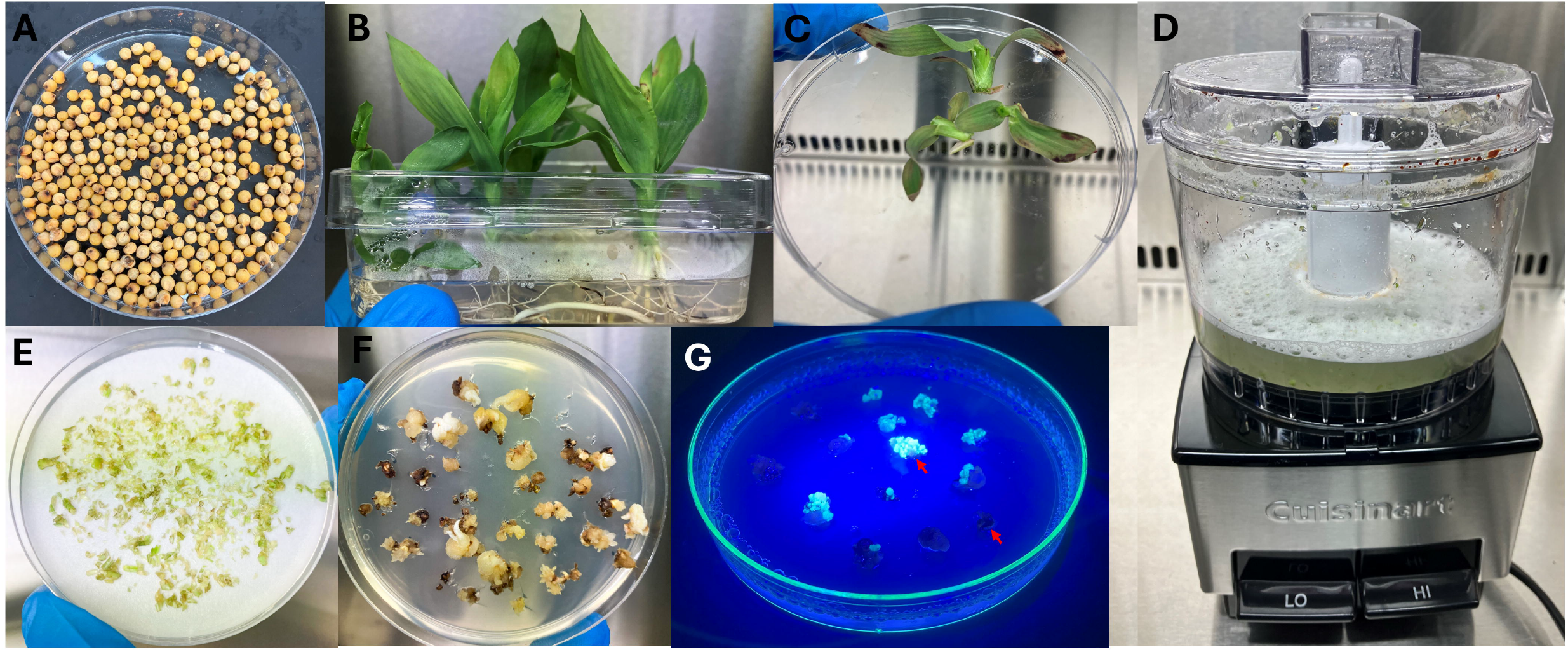
Sorghum leaf transformation following methods from Wang *et al*., 2023 and transgenic embryonic calli detection using AmCyan. **A)** Sorghum seeds were surface sterilized using chlorine gas and selected based on size and shape for planting. **B)** Sorghum seedlings grown for twenty-one days in tissue culture were used for explant source material. **C)** Leaf base segments were prepared from cutting seedlings ∼2 cm abaxial to mesocotyl, removing outer leaves, and cutting longitudinally (not shown). **D)** Prepared leaf base tissue was placed in a mini blender with an Agrobacterium suspension (90 ml) and blended 10x times using short pulses. **E)** Prepared leaf explants were collected through a metal sieve and spread on filter paper covering co-cultivation media (710N). **F)** After three days co-cultivation, explants were transferred to resting media (13266P) for four-weeks to assess survival and **G)** visualized for AmCyan detection at two-weeks. AmCyan signal from transgenic calli was scored as either positive (top red arrow) or negative (bottom red arrow) and is shown in example without excitation filter to emphasize signal differences.

### Sorghum leaf-base transformation using altruistic T-DNA vectors

The application of altruistic vectors has been effective for improving monocot transformation by separating morphogenic regulators and target cargo on separate T-DNAs and optimizing ratios of the different Agrobacterium strains to capture transgenic events (Hoerster *et al*., 2020; Mookkan *et al*., 2017; Kang *et al*., 2023). This approach can eliminate the need for removal of morphogenic regulators either during or post-transformation and simplify target vector construction.

To test this approach, a traditional altruistic T-DNA vector was constructed using GRF4-GIF1 (A-GRFGIF, **Figure 1B**) –the same backbone as C-GRFGIF – and the T-AmCyan vector as cargo (**Figure 3**). Consistent with previous results using traditional vectors, transformation experiments with altruistic vectors resulted in both non-viable (**Figure 3B**) and viable embryonic calli viable (**Figure 3C**) with viable embryonic calli capable of regeneration to whole plants (**Figure 3E**). However, the average percentage of viable calli was higher using the altruistic GRF4-GIF1 vector (61.3 ± 4.7%, **Table 2**) compared to the traditional GRF4-GIF1 vector (43.2 ± 2.9%, **Table 1)** and resulted in an average of 21% more calli positive for AmCyan. These results suggest altruistic delivery of morphogenic regulators can improve monocot leaf-base transformation and supports previous studies using conventional explant types.

**Table 2.**
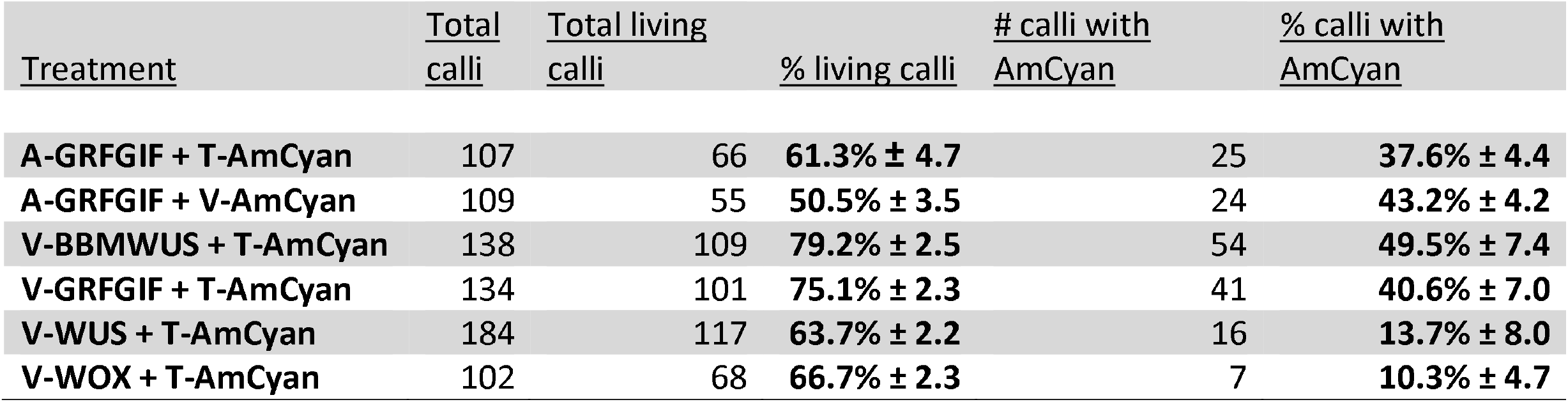
Transgenic embryonic calli efficiency using altruistic FoMV-mediated sorghum leaf transformation. Data was collected using twenty seedlings per treatment. Total embryonic calli was counted two weeks post infection (“total calli”). Calli survival was assessed by counting calli capable of growth at four weeks post infection (“total living calli”). Calli transgenesis was assessed using fluorescent AmCyan imaging two weeks post infection. Calli survival was calculated as a percentage (“% calli survival”) of “total living calli” divided by “total calli.” Calli transgenesis was calculated as a percentage by dividing the number of AmCyan positive calli (“# calli with AmCyan”) by “total living calli.” Average percentages of five to eight biological replicate plates are provided with standard errors (**Table S2**).

| <u>Treatment</u> | <u>Total</u><br><u>calli</u> | <u>Total living</u><br><u>calli</u> | <u>% living calli</u> | <u># calli with</u><br><u>AmCyan</u> | <u>% calli with</u><br><u>AmCyan</u> |
| --- | --- | --- | --- | --- | --- |
| **A-GRFGIF + T-AmCyan** | 107 | 66 | **61.3% ± 4.7** | 25 | **37.6% ± 4.4** |
| **A-GRFGIF + V-AmCyan** | 109 | 55 | **50.5% ± 3.5** | 24 | **43.2% ± 4.2** |
| **V-BBMWUS + T-AmCyan** | 138 | 109 | **79.2% ± 2.5** | 54 | **49.5% ± 7.4** |
| **V-GRFGIF + T-AmCyan** | 134 | 101 | **75.1% ± 2.3** | 41 | **40.6% ± 7.0** |
| **V-WUS + T-AmCyan** | 184 | 117 | **63.7% ± 2.2** | 16 | **13.7% ± 8.0** |
| **V-WOX + T-AmCyan** | 102 | 68 | **66.7% ± 2.3** | 7 | **10.3% ± 4.7** |

**Figure 3.**
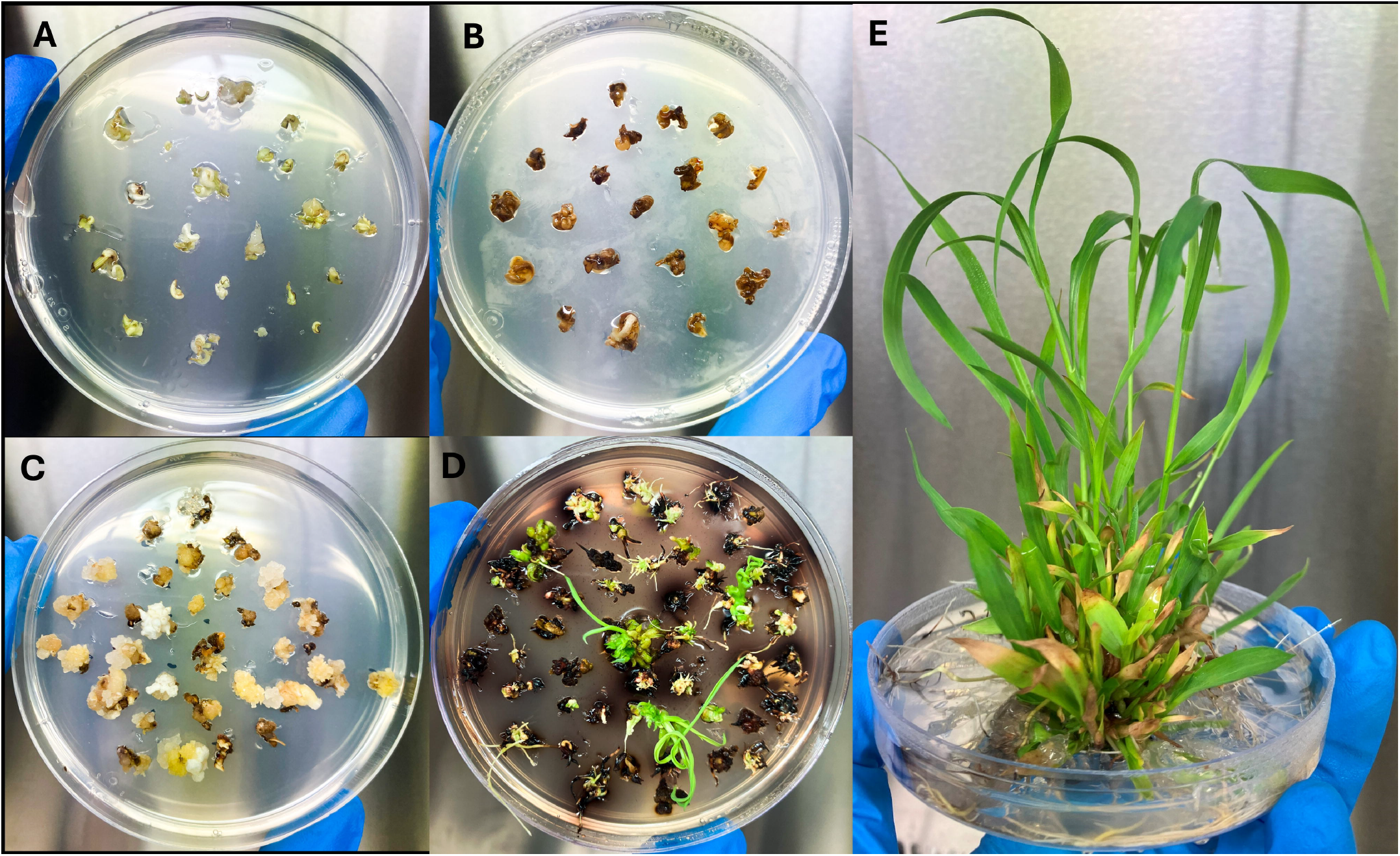
Sorghum regeneration from leaf explants using conventional and altruistic T-DNA vectors and methods adapted from Wang *et al*., 2023. **A)** Two-week-old embryonic calli generated on resting media (13266P) following Agrobacterium co-culture (example: C-GRFGIF). Four-week-old embryonic calli grown on resting media with **B)** no calli survival (example: C-GRFGIF) and **C)** ∼50% calli survival (example: A-GRFGIF with T-AmCyan). **D)** Regenerated plantlets following two weeks on shooting media (13329A) (example: A-GRFGIF with T-AmCyan). **E)** Rooted regenerated plantlets following four-weeks on rooting media (13158) (example: A-GRFGIF with T-AmCyan).

### Viral-mediated transgene expression in sorghum leaf-base explants

Viral-based vector systems are capable of many of the expression requirements for morphogenic regulators in transformation protocols. Positive-strand, RNA viruses, such as FoMV have been adapted for delivery by T-DNA vector systems, have mild symptoms, and have been modified to incorporate sub-genomic promoters from other viruses to enable high, transient transgene expression without relying on direct T-DNA delivery (Mei *et al*., 2019; Mei *et al*., 2016).

To test application of viral vectors for leaf transformation, a previously developed FoMV vector system was used to incorporate sequences either for AmCyan (V-AmCyan), as a negative control and for viral tracking, or morphogenic regulators (V-MR) (**Figure 4**). Four different morphogenic regulator combinations were cloned into FoMV and all incorporated sequences were driven by the same Pea Early-Browning virus (PeBV) sub-genomic promoter. Before morphogenic regulators could be tested, it first needed to be determined if the virus was capable of replication and did not impair embryonic calli formation. To this end, V-AmCyan was delivered in combination with the traditional altruistic vector, A-GRFGIF, and observations of embryonic calli formation and AmCyan detection were made post infection.

**Figure 4.**
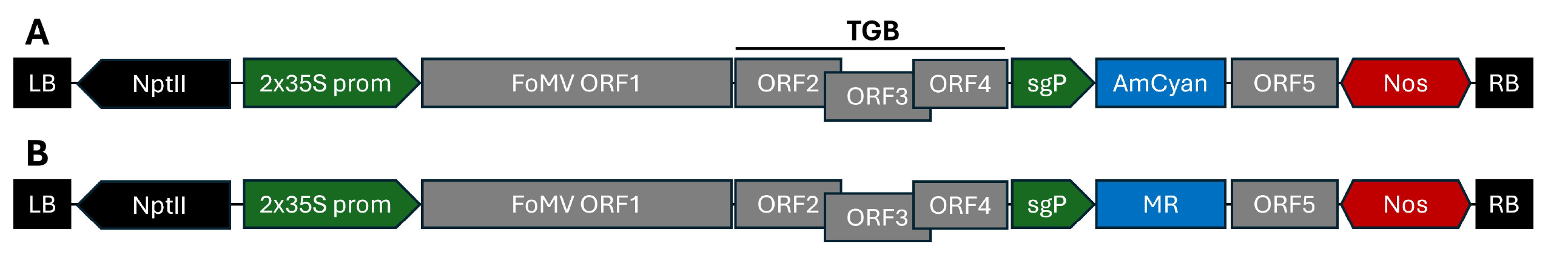
Foxtail Mosaic Virus (FoMV) T-DNAs used for sorghum leaf transformation. Control (**A** [V-AmCyan]) and altruistic (**B** [V-MR]) vectors for viral tracking and altruistic virus-mediated transformation, respectively. Morphogenic regulators used in V-MR vectors include maize *Wus2* (V-WUS), maize *Wox2a* (V-WOX), maize *Bbm*/*Wus2* (V-BBMWUS) and wheat GRF4-GIF1 with a P2A translational skipping signal used for expressing maize *Bbm*/*Wus2* (not shown). Green arrows indicate promoter or sub-genomic promoter (sgP) elements. Gray boxes indicate consecutive and overlapping FoMV open reading frames (ORFs), with ORFs 2, 3 and 4 comprising the triple gene block (TGB). Blue boxes indicate coding sequences. Red hexagons indicate terminator elements. Black boxes indicate right and left T-DNA borders. Black arrows indicate selection marker cassettes driven by the 2x 35S promoter and 35S terminator. Coding sequences were incorporated using Gibson assembly and SapI restriction enzyme sites.

AmCyan detection in leaf-base explants post infection in all treatments was complicated by phenolic production and residual Agrobacterium on explants. Nevertheless, leaf-base explants infected with V-AmCyan resulted in average percentages of both viable embryonic calli and embryonic calli with AmCyan at four and two-weeks post infection, respectively, similar to altruistic delivery experiments incorporating T-AmCyan (**Table 2**). However, unlike T-AmCyan treatments, AmCyan signal in V-AmCyan treatments were largely diminished at four-weeks post infection.

To verify these results, total RNA was extracted from embryonic calli at two and four-weeks post infection and subjected to reverse-transcriptase PCR (RT-PCR) for viral RNA detection (**Figure S2**). Whereas viral RNA was detected at two-weeks post infection, only residual levels were detected at four-weeks. These results reflect observations of AmCyan in embryonic calli and suggest viral replication is transient in leaf-base explant and embryonic calli tissues.

### Viral-mediated leaf-base transformation and regeneration in sorghum

Previous experiments incorporating AmCyan into viral vectors demonstrated viral replication and transient transgene expression in leaf-base transformation tissues (Zhong *et al*., 2024; Baysal *et al*., 2024b). However, AmCyan is a relatively small transgene (690 bp) and most morphogenic regulators are much larger. Due to FoMV and other positive-strand RNA viruses having cargo-capacity limits of approximately 2,000 bp (Mei *et al*., 2016; Liu *et al*., 2016; Avesani *et al*., 2007), it was unclear if viral expression of morphogenic regulators was sufficient for leaf-base transformation.

To accommodate these restrictions, maize *Wox2a* (975 bp: V-WOX) and *Wus2* (909 bp: V-WUS) were tested alongside GRF4-GIF1 (1,911 bp: V-GRFGIF) and *Bbm*/*Wus2* (2,130 bp: V-BBMWUS) as altruistic viral vectors for delivery of T-AmCyan. Interestingly, V-BBMWUS outperformed all other altruistic treatments for both embryonic calli formation and embryonic calli with AmCyan, with V-GRFGIF producing similar results (**Table 2**). Furthermore, both V-WUS and V-WOX generated similar levels of embryonic calli compared to A-GRFGIF but lower levels of AmCyan positive embryonic calli. Similar to the results of V-AmCyan, only residual levels of virus were detected in embryonic calli of morphogenic regulator treatments at four-weeks post infection, supporting transient expression of morphogenic regulators from viral vectors (**Figure S2**).

Previous studies have shown persistent expression of morphogenic regulators can impair regeneration of embryonic calli to shoots (Gordon-Kamm *et al*., 2019; Lowe *et al*., 2016). Hence, transient expression of morphogenic regulators in viral delivery experiments might allow shoot formation without the need for recombinase-mediated excision. To test this, embryonic calli from V-MR altruistic delivery experiments were observed for shoot formation at four and six-weeks post infection. Differing from previous experiments using traditional vectors, immature shoots with AmCyan signal could be observed at four-weeks in V-MR treatments as opposed to six to eight-weeks (**Figure 5A-B**). These AmCyan positive shoots continued to develop into regenerated shoots (**Figure 5C-D**). To validate AmCyan transgene delivery, total RNA and genomic DNA was extracted from embryonic calli and mature shoots and subjected to RT-PCR and PCR analysis, respectively (**Figure 5E-F**). AmCyan mRNA in AmCyan-positive calli and regenerated shoots was clearly detected with some background in AmCyan-negative calli, likely from residual Agrobacterium. Furthermore, the AmCyan transgene was most clearly detected in AmCyan-positive, regenerated shoots with some detection in negative calli and background signal in wild-type controls. Collectively, these results demonstrate the utility of viral expression of morphogenic regulators for leaf-based transformation that could be applied to other transformation protocols.

**Figure 5.**
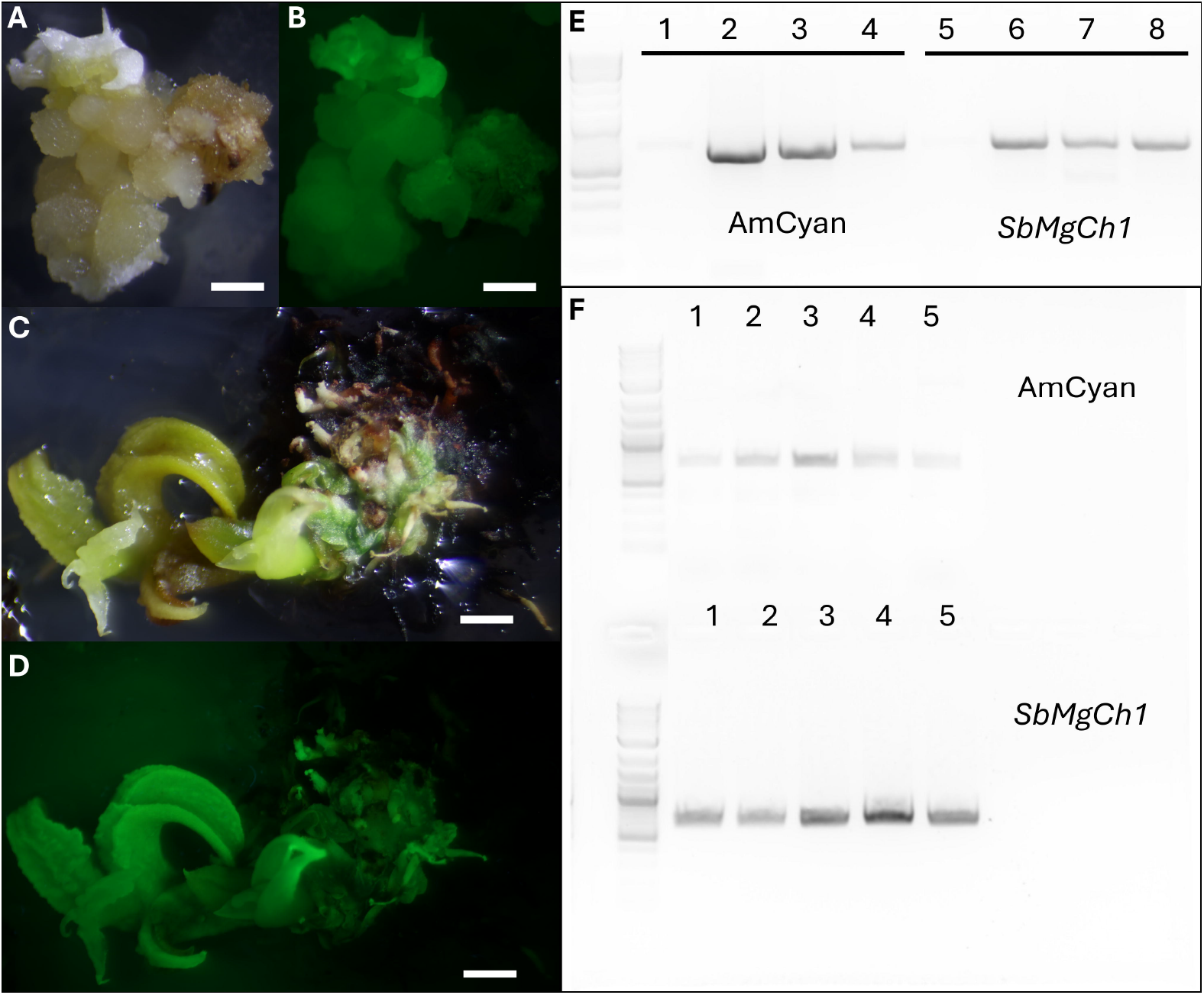
Sorghum regeneration from leaf explants using altruistic FoMV T-DNA vectors and methods adapted from Wang *et al*., 2023. Four-week-old embryonic calli regenerated on resting media shown using light (**A)** and fluorescent **(B)** imaging. Example: V-GRFGIF + T-AmCyan. Six-week-old shoots regenerated on shooting media shown using light **(C)** and fluorescent **(D)** imaging. Example: V-WOX + T-AmCyan. **E)** RT-PCR validation of the AmCyan transgene in sorghum embryonic calli and regenerated shoots using water control (lanes 1, 5), regenerated shoot (lanes 2, 6), AmCyan positive calli (lanes 3, 7) and AmCyan negative calli (lanes 4, 8) total RNA templates, and AmCyan (lanes 1-4) and *SbMgCh1* (lanes 5-8) primers. **F)** PCR validation of the AmCyan transgene in sorghum embryonic calli and regenerated shoots using AmCyan negative calli (lane 1), AmCyan positive calli (lane 2), regenerated shoot (lane 3), and Tx430 wild-type (lanes 4, 5) genomic DNA templates, and AmCyan (top lanes) and *SbMgCh1* (bottom lanes) primers. Calli and shoots sampled at two and six-weeks for RT-PCR and PCR, respectively. Example: V-WOX + T-AmCyan. Scale bar: 5mm. DNA marker: 2-log DNA ladder.

## Discussion

Transformation of monocot species remains a critical bottleneck to employing modern genetic improvement tools and conducting functional genomics (Altpeter et al., 2016; Parrott et al., 2024; Butler et al., 2017; Cardi et al., 2023). Important progress has been made in monocot transformation with the incorporation of morphogenic regulators, first demonstrated in maize using immature embryos and a combination of *Bbm* and *Wus2* (Lowe *et al*., 2016). This approach was quickly extended to sorghum transformation and has eventually incorporated additional morphogenic regulators, such as *Wox5*, GRF5, and GRF4/GIF1 leading to improved protocols in other cereals, such as wheat, rye, barley, triticale and rice (Youngstrom *et al*., 2024; Gordon-Kamm *et al*., 2019). Nevertheless, these protocols rely on immature embryo explants, which are labor-intensive and difficult to generate.

Leaf explants derived from leaf bases have historically been shown to be effective for generating callus in a range of grass species, including rice (12.44% callus/plant) (Tran and Sanan-Mishra, 2015), maize (65% callus/explant) (Ahmadabadi *et al*., 2007) and ma bamboo (53% callus/explant) (Ye *et al*., 2017). However, efficiency of regenerable embryonic callus production in these species has either been low or non-transgenic. For these reasons, more recent advancements in leaf base transformation, including those conducted by Wang *et al*., 2023, have demonstrated the importance of vector design by use of the 3xEnh-Ubi promoter driving *Bbm* expression and a broader range of ubiquitous monocot promoters driving *Wus2* expression. These optimizations resulted in embryonic callus generation ranging from 50-80%. However, these efficiencies depend on combining morphogenic regulator gene cassettes with target gene cassettes on delivered T-DNAs. This results in large, complex vectors that can be difficult to assemble and require Cre/lox excision of morphogenic regulator genes to regenerate shoots.

Plant RNA viruses, such as FoMV and BSMV have been demonstrated to be effective for virus-mediated expression of transgenes for transient protein expression in both monocots and dicots (Mei et al., 2019; Baysal et al., 2024; Bedoya et al., 2010). The transient nature of virus-mediated gene expression is thought to be the result of host plant defenses and potential viral genome instability due to heterologous sequence incorporation (Abrahamian *et al*., 2020; Scholthof and Scholthof, 2023; Khakhar and Voytas, 2021). Viral genome stability is a particularly important factor for positive-strand RNA viruses which have more strict cargo capacities compared to negative-strand RNA viruses. For example, a PVX expression system showed an increased ratio of wild-type (no-insert) vector to recombinant (insert) vector with increasing insert sizes, ranging from 200 bp to 1,700 bp, demonstrating the effects of cargo capacity on virus stability (Avesani *et al*., 2007). Furthermore, this study and others have shown similar viral instability in subsequent passages from plants that were initially infected, indicating viral-mediated expression of transgenes can be transient under certain conditions (Whitham *et al*., 2007; Khakhar and Voytas, 2021).

The first objective of this study was to reduce the complexity of vectors required to induce embryonic callus formation in monocot leaf transformation protocols using sorghum transformation as a proof-of-concept. Co-delivery of altruistic vectors carrying morphogenic regulators with target T-DNAs has been an effective approach in protocols using immature embryo explants (Hoerster *et al*., 2020; Mookkan *et al*., 2017; Kang *et al*., 2023) and was tested in this study using GRF4/GIF1. Consistent with these previous studies, altruistic delivery resulted in comparable efficiencies of embryonic callus generation compared to conventional leaf transformation vectors, regardless of morphogenic growth regulator, and was outperformed only by the conventional *Bbm*/*Wus2* vector for generation of transgenic embryonic callus. This provides valuable information towards future leaf transformation vector design, suggesting an altruistic approach can be used incorporating a single altruistic vector carrying morphogenic regulator genes in combination with different target vectors carrying genes of interest. This approach not only reduces complexity of vector design and construction but could also reduce morphogenic regulator gene integration and linkage between regulator genes and target genes.

The second objective of this study was to reduce protocol complexity by eliminating recombinase-mediated excision of morphogenic regulator genes and the need for inducible recombinase expression. Excision of morphogenic regulator genes via inducible recombinase expression is an important step in transformation protocols not only to eliminate unwanted effects of morphogenic regulators in downstream transformation steps and resulting events, but also to reduce toxicity associated with high recombinase expression (Wang *et al*., 2023; Gordon-Kamm *et al*., 2019; Wang *et al*., 2020). To this end, morphogenic regulator genes ranging from 909 bp to 2,130 bp in size were cloned into the FoMV expression vector and delivered altruistically to sorghum leaf explants. Surprisingly, the largest morphogenic regulator genes (*Bbm*/*Wus2*, GRF4-GIF1) outperformed smaller regulator genes (*Wus2, Wox2a*) and conventional altruistic vectors for both formation of total embryonic and transgenic callus. Regardless of size, virus detection in both embryonic calli and shoots was drastically reduced after four weeks post infection, suggesting morphogenic regulator genes were expressed transiently at early stages of callus formation.

Modern tools for genetic improvement of plants are continuously improving with great strides being made in recent years with the utilization of morphogenic growth regulators and plant RNA viruses. This study demonstrated that these technologies can be combined to increase efficiencies of embryonic calli formation, a limiting step in monocot leaf transformation, to improve protocol efficiencies and reduce experimental complexity. This approach can be further improved with incorporation of alternative plant virus expression systems, such as TRV and BSMV which could be combined or used individually to expand application to additional dicot and monocot species and improve transformation efficiencies. Furthermore, alternative viral promoters could be used to optimize morphogenic regulator gene expression or morphogenic regulator gene coding sequences optimized for expression in viral systems. Nevertheless, the continued improvement of leaf transformation protocols in monocot species is promising and has the potential to have important impacts on crop production worldwide.

## Experimental procedures

### Vector construction

Traditional T-DNA vectors were constructed using components and methods from Chamness et al., 2023. The maize HSP (*Hsp17*) promoter and triple enhancer (3xEnh) sequences were taken from Wang *et al*., 2023 and cloned into “Level 0” modules using Golden Gate cloning and BbsI sites.

Viral T-DNA vectors were constructed using Gibson assembly and the FoMV cloning vector, pEE082 linearized using SapI (Baysal *et al*., 2024). PCR reactions for generating assembly fragments were performed using Chamness *et al*., 2023 components as templates, oligos listed in **Table S1**, and the Q5 DNA polymerase (New England Biolabs). Gibson assembly was conducted using the Gibson Assembly® Master Mix (New England Biolabs) with recommended conditions and 75 ng linearized vector.

### Plant materials

*Sorghum bicolor* cv Tx430 seed used for sorghum leaf-base transformation experiments were generated by growing plants in a greenhouse at 28°C and a photoperiod of 16/8 h day/night. Sorghum seed was sterilized using chlorine gas for 16 h and germinated in Phytatrays™ (Sigma-Aldrich) in a growth incubator at 28°C with a photoperiod of 16/8 h day/night. Twenty-four to twenty-six seedlings were used per treatment in transformation experiments.

### Sorghum leaf-base transformation

Sorghum leaf-base transformation was carried out following methods from Wang *et al*., 2023 with the following modifications. Fourteen to twenty-one-day-old sorghum seedlings were used for starting explant material and embryonic calli were transferred to 13266P resting media without filter paper. No selection or heat shock treatments were used in any treatments.

### Evaluation of embryonic calli viability and detection of AmCyan fluorescence

Embryonic calli generated in sorghum leaf-base transformation experiments began to emerge approximately seven to ten days after Agrobacterium infection. At fourteen days post infection, embryonic calli were transferred to 13266P resting media, counted (“total calli”), and visualized for AmCyan fluorescence (“# calli with AmCyan”). Embryonic calli were counted as being positive for AmCyan only if high intensity fluorescence was observed (**Figure 2**). Embryonic calli were allowed to grow for an additional fourteen days, or four weeks post infection in the dark before surviving calli were counted (“total living calli”). Embryonic calli were considered viable if dividing, friable calli and somatic embryos could be identified. AmCyan detection was carried out using the Xite Fluorescence Flashligh System (Electron Microscopy Sciences, PA, USA) with a Royal Blue, single-wavelength excitation flashlight (440-460 nm) and detected using a 500-560 nm bandpass filter. Images were taken with a Canon EOS 80D camera with a Canon EF-S 18-55 mm f/3.5-5.6 IS STM lens (Canon USA, Lake Success, NY, USA).

### RT-PCR and PCR analysis

Total RNA for RT-PCR analysis was extracted using the RNeasy Plant Mini Kit following recommended conditions (Qiagen, Germantown, USA). 100 ng total RNA was used for RT-PCR using the OneStep RT-PCR Kit, recommended conditions for 25 μL reactions, and oligos listed in **Table S1**. Cycling conditions for all RT-PCR experiments included a one-minute 72C extension and thirty-five cycles. For detection of AmCyan mRNA, a 66°C annealing temperature was used. For detection of the FoMV TGB2, a 63°C annealing temperature was used. The sorghum *Magnesium-chelatase subunit I* (*SbMgCh1*) mRNA was used as a reference and used a 63°C annealing temperature.

Total genomic DNA for PCR analysis was extracted using the DNeasy Plant Pro Kit following recommended conditions (Qiagen, Germantown, USA). 150 ng total genomic DNA was used for PCR using the Q5 DNA polymerase (New England Biolabs), recommended conditions for 50 μL reactions, and oligos listed in **Table S1**. Cycling conditions for all PCR experiments included a 66°C annealing temperature, one-minute 72C extension and thirty-two cycles.

## Author contributions

N.M.B. conceived the research and conducted the experiments. A.T.C. and C.S. assisted in transformation experiments and vector cloning, respectively. N.M.B. and D.F.V. wrote the manuscript.

## Acknowledgements

We thank Albert Kausch and his lab for assistance in adapting and obtaining materials related to the sorghum leaf transformation protocol, and Steve Whitham for providing the FoMV viral vectors. Funding for this work was provided by the Department of Energy Biological and Environmental Research (BER) Program (US Department of Energy, Award Number DE-SC0023160).

## Supporting information

**Figure S1**. Sorghum leaf-base transformation using control traditional and viral T-DNA vectors

**Figure S2**. RT-PCR detection of FoMV in sorghum embryonic calli

**Table S1**. Oligos for FoMV cloning, RT-PCR and PCR analysis

**Table S2**. Data set for Table 2

