## Supplementary figures and images for "Viral-mediated delivery of morphogenic regulators enables leaf transformation in *Sorghum bicolor* (L.)"

### supplemental figures

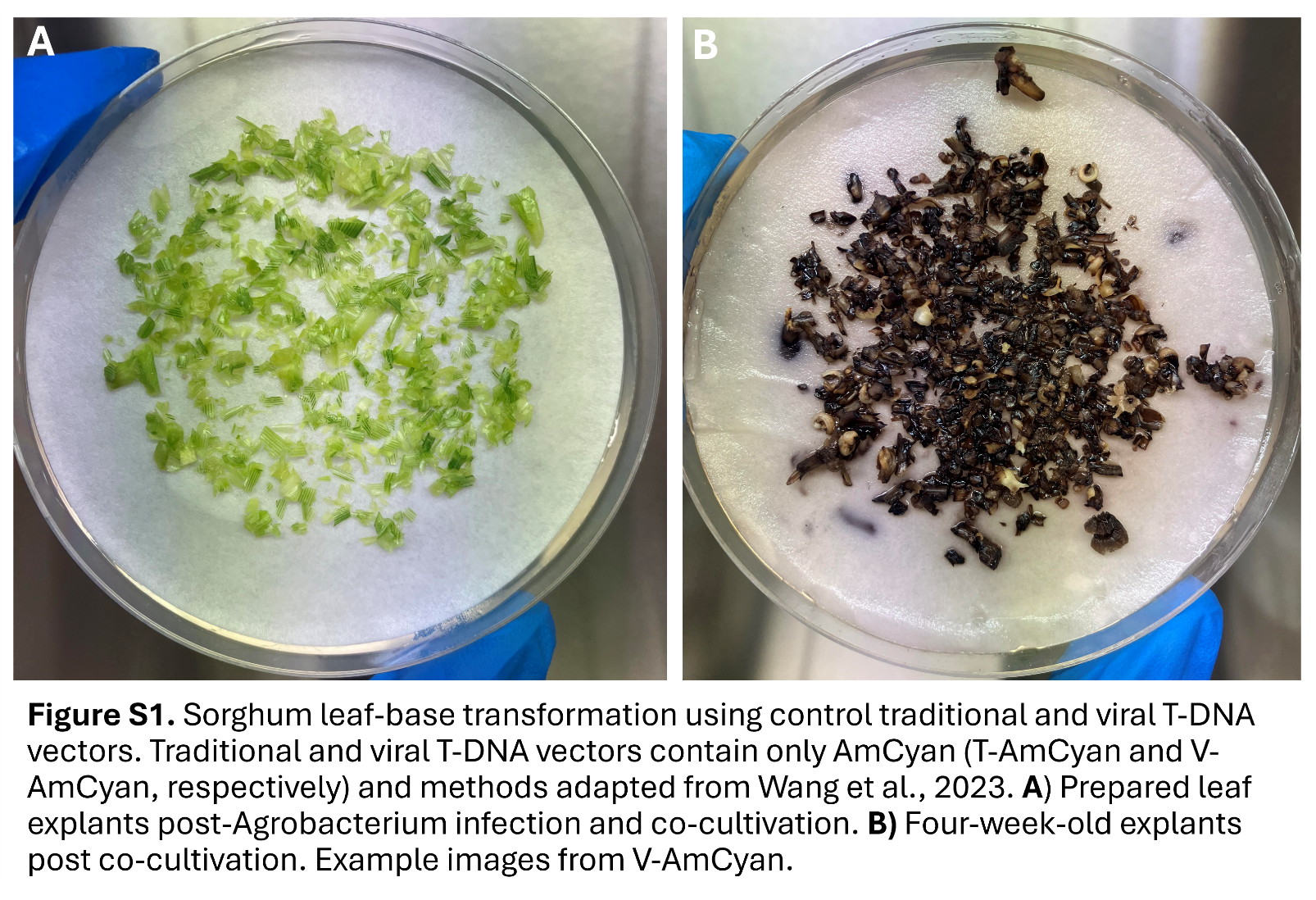

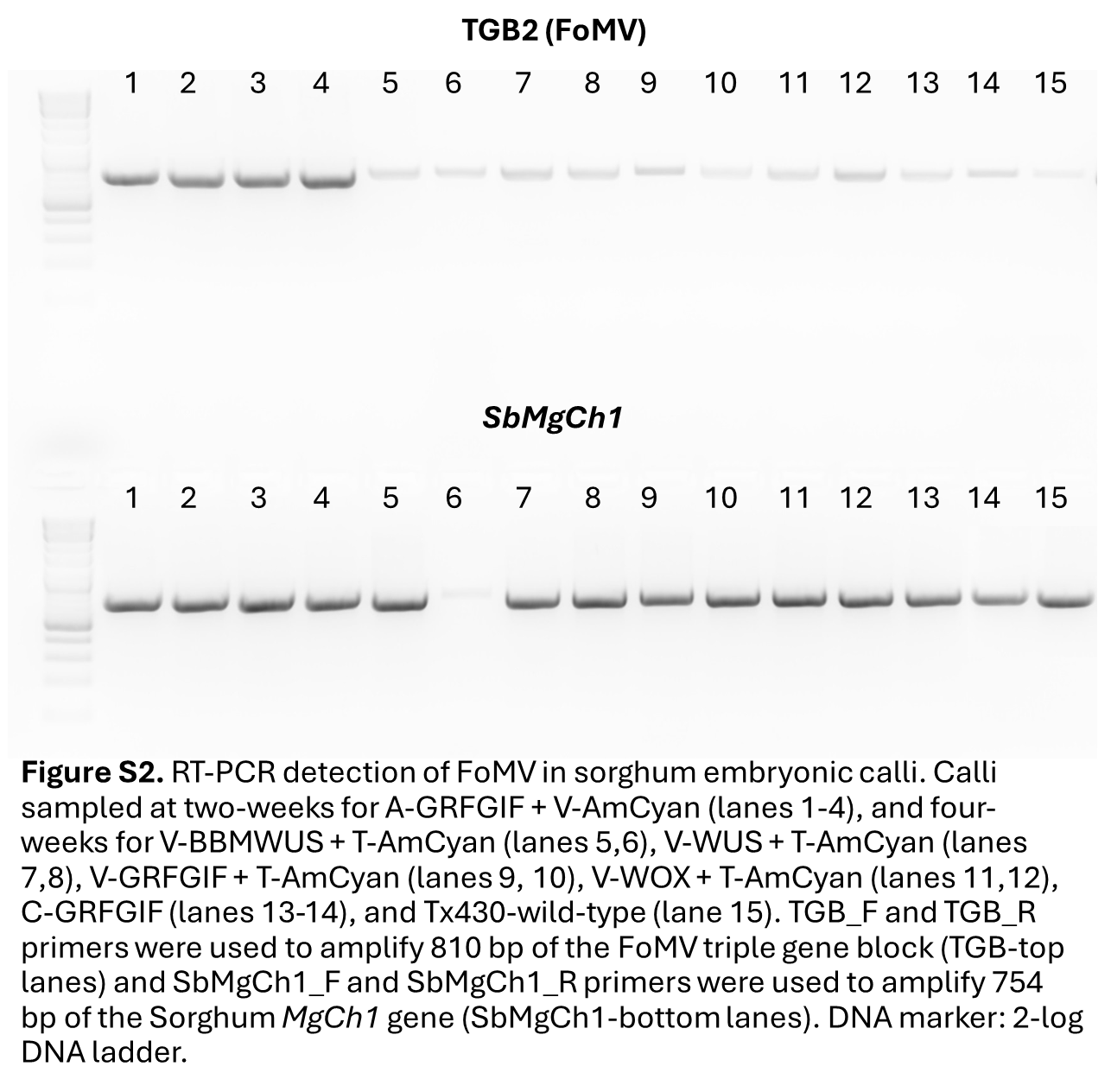
